# Contact calling is predicted by cooperative relationships in vampire bats

**DOI:** 10.64898/2026.03.04.709596

**Authors:** Julia K. Vrtilek, Haley J. Gmutza, Sydney Decker, Gerald G. Carter

## Abstract

Group-living animals often coordinate their behavior using “contact calls”. Identifying the function of these calls requires testing whether they are intended for any group member or targeted to specific preferred associates. If contact calling is used to coordinate with preferred associates, then higher rates of contact calling are expected between group members with a history of more frequent affiliation and cooperation. To test this hypothesis, we constructed a contact-calling network using synchronized recordings of vocal interactions between all 28 possible pairs of 8 female common vampire bats with well-sampled histories of social grooming and regurgitated food sharing. Bayesian multilevel models show that pairwise rates of contact calling were clearly predicted by social grooming and cooperative allofeeding rates in ways not explained by kinship. These findings show that common vampire bats use contact calls to coordinate with specific same-sex associates, unlike other studied bat species where individuals produce contact calls at similar rates towards different group members. We also found that, compared to white-winged vampire bats, common vampire bats are ten times less likely to rapidly respond to a contact call; this suggests yet-to-be-discovered differences in social behavior between vampire bat species. Finally, we discuss implications for the ‘vocal grooming’ hypothesis.

## INTRODUCTION

Many group-living animals use vocalizations to coordinate movements with group members or specific social partners (1). To understand the function of these “contact calls”, it is necessary to identify the intended receivers, which could be any group member (2) or a specific individual such as a mate (3,4), close affiliate (5–8), or one’s offspring (9)). However, identifying intended receivers is difficult in a naturalistic group setting. One potential solution is to record contact calling among isolated pairs of animals, then use multi-level models, such as the social relations model (10), to disentangle how well variation in calling rates are explained by caller, receiver, and their unique caller-receiver relationship. In this scenario, if calls are addressed to a group, variation in calling rates would be explained by the caller but not the receiver, as e.g. in disc-winged bats (2). Alternatively, if calls are addressed to specific receivers, rates of calling should also depend on the caller-receiver relationship. Furthermore, if callers use contact calls to coordinate or cooperate with preferred associates, we expect higher rates of contact calling between group members with a history of more frequent affiliation and cooperation.

To test this hypothesis, we constructed a contact-calling network using synchronized recordings of vocal interactions between pairs of female common vampire bats (*Desmodus rotundus*) with known histories of social grooming (allogrooming) and regurgitated food sharing (allofeeding). Unrelated female vampire bats use reciprocal allogrooming to develop reciprocal allofeeding relationships (11), and these cooperative relationships can prevent starvation when bats fail to feed (12). When female vampire bats are isolated from their colony, they produce contact calls that allow preferred associates to identify and find them (13). Contact call convergence occurs among familiar but genetically unrelated vampire bats, with the greatest convergence between food-sharing partners (14). However, it remains unclear whether vampire bats produce or exchange more contact calls with preferred cooperation partners.

Here, we show that contact-calling rates within vampire bat pairs are clearly predicted by the pair’s allogrooming and allofeeding in ways that are not explained by kinship. Because both allogrooming and allofeeding are highly reciprocal (11,15), we also assessed whether contact-calling networks were similarly reciprocal. To do this, we first fit a social relations model allowing us to estimate the correlation between calls given and calls received, either by the current partner (dyadic vocal reciprocity) or by all partners (generalized vocal reciprocity). To explain why call rates might be reciprocal, we then measured how often common vampire bats respond to each other in rapid vocal exchanges.

## METHODS

### Estimating genetic relatedness and social networks

Subjects were eight individually tagged adult female common vampire bats, originating from the Cincinnati Zoo and kept in captivity at The Ohio State University. To estimate genetic kinship among all 22 vampire bats in this colony, we first collected 3-mm wing punch tissue samples (stored in 95% ethanol) and extracted DNA using the Qiagen DNeasy blood and tissue kit. We quantified extracted DNA using a Qubit 4 fluorometer and prepared samples for Genotyping-by-Sequencing (GBS) following Elshire et al. protocol (16) and using the restriction enzyme PstI. The GBS library was sequenced at The Ohio State University Comprehensive Cancer Center Shared Genomics Research Laboratory on an Illumina NextSeq 2000, sequencing 100bp single-end reads. We assessed sequence read quality with FastQC v0.11.8 (17). We used the iPyrad pipeline (18) to demultiplex samples, trim adapters, and map reads using the Bat1K *Desmodus rotundus* reference genome (NCBI GCF_022682495.1) (19). We filtered reads to keep only loci that are mapped to autosomal chromosomes using PLINK2 (20). To estimate genetic relatedness matrices from our genomic datasets, we used the KING-robust estimator (21).

We estimated rates of allogrooming and allofeeding across all bats. To induce allofeeding and allogrooming, we simulated a focal bat’s failure to feed overnight by isolating and fasting each bat on 4 different nights, following the procedure from Carter & Wilkinson (15). Briefly, each focal bat was fasted for 25-33 hours, while all other bats were fed to allow for regurgitated food donations. The fasted bat was then placed back into the flight cage with all other bats and focal-followed for one hour using a Sony Nightshot HDD 10.2 Megapixel video camera and infrared spotlight. We measured the duration of allogrooming events (one bat licking another bat for at least 5 seconds) and allofeeding events (one bat licking the mouth of another for longer than 5 seconds) following Carter & Wilkinson (15).

### Estimating the contact calling network

To record pairwise vocal interactions, we isolated pairs of bats such that they could hear each other but not other bats in their colony (see Fig. S1). We placed two plastic bins (∼100 liters in volume) about 15 centimeters apart, each lined with acoustic foam to dampen echoes and increase recording quality. Within each bin, a bat was placed inside a tube of plastic mesh to constrain it within 10-30 cm of an Avisoft CM16 ultrasound condenser microphone (frequency range 10-200kHz, Avisoft Bioacoustics, Berlin, Germany). The highly directional microphones faced directly away from each other, allowing us to unambiguously record only the intended bat. Sound was digitized with 16-bit resolution at a sampling rate of 500 kHz through a four-channel Avisoft UltraSoundGate 416H to a laptop running the program Avisoft Recorder. To confirm that we were recording sounds of the same amplitude equally at both microphones, we calibrated them using the same playback of bat calls from an Avisoft UltraSoundGate Player BL Pro. We recorded all possible pairwise combinations of 8 bats (28 pairs) for an hour at a time, and repeated each one-hour recording session three times over the course of 23 days, for a total of 3 recorded hours per pair and 84 total hours recorded.

We identified contact calls from WAV files using methods described in Vrtilek et al. (14). To filter out other sounds, we excluded call selections with peak frequencies below 10 kHz and durations less than 3 ms or longer than 50 ms. We also excluded possible sounds which did not have a clear fundamental frequency syllable. Finally, we manually checked spectrograms of all selections and excluded 49 sounds that were obviously not bat calls. Although one bat never called, we recorded 3687 total contact calls from the others (mean calls per caller = 527, minimum = 24, maximum = 2038 calls).

### Caller-receiver models

We estimated the effect of social affiliation as a predictor of contact-call counts from each bat across 3 recording sessions. To do this, we fit a Bayesian negative binomial multi-level model using the R package *brms* (22), with caller and receiver as crossed random effects (grouping variables with varying intercepts) and one fixed effect indicating social preference (either allogrooming rate towards the recipient, allofeeding rate towards the recipient, or genetic kinship with the recipient). Each fixed effect was scaled (z-transformed). We did not include all three predictors in one model because they are redundant proxies of the same underlying effect (i.e. latent social preferences).

We call this model the *“caller-receiver model”* because it assumes calling rates are caused by the caller (some bats call more than others), the receiver (some bats receive more than others), and a caller-receiver relationship or social preference, which is captured by the fixed effect. We used leave-one-out cross validation to compare these models to ones that included dyad as an additional multi-membership random effect; we found that this additional grouping variable did not significantly change the results or increase out-of-sample predictive accuracy in any of the models, so we present the simpler model to reduce potential confounding.

We used default priors. We used warmup and chain lengths of 2000 samples. To ensure Markov Chain Monte Carlo (MCMC) convergence, we checked that all chains reached the same estimates for coefficients (minimum and maximum Rhat = 1.00). We used posterior predictive checks to assess model fit.

### Social relations models

For reasons described further in the discussion, we also modelled calling rates with a more complex Bayesian social relations model (Poisson distribution with dyad-level random effects) using the R package *STRAND* (10). We again used default priors, warmup and chain lengths of 2000 samples each, and confirmed chain convergence (minimum and maximum Rhat = 1.00).

Compared to the *caller-receiver model*, the *social relations model* makes two different key assumptions. First, by incorporating a dyad-level random effect, it assumes that bats can have preferred contact calling relationships that are not fully captured by the social preferences estimated by allogrooming, allofeeding, or kinship. Second, it assumes that the random effects of caller and receiver are correlated at both the dyad-level (calls from bat A to B predict calls from bat B to A) and the group-level (calls given by bat A to all receivers predict calls received by bat A from all callers) for reasons beyond the social preferences estimated by allogrooming, allofeeding, or kinship.

Finally, to contextualize the dyadic effects and dyad-level correlation for contact calling, we compared a social relations model of contact calling predicted by kinship to a similar social relations model with allogrooming predicted by kinship. We used the same priors and convergence checks (minimum and maximum Rhat = 1.00).

### Testing for rapid vocal exchanges

To test for rapid vocal exchanges, we first measured the lag time between alternating calls from different callers as the time between the end of the first call and the start of the second. Next, we defined antiphonal calls as those within 0.5 s of a partner’s contact call, and calculated the frequency of antiphonal calling as the number of antiphonal contact calls divided by the total number of recorded contact calls. We used this measure to allow for direct comparison with the same measure reported from white-winged vampire bats (23). We also calculated the frequency of a bat getting an antiphonal response to a call (the number of antiphonal calls each bat received divided by the number of calls each bat made) and the frequency of a bat giving an antiphonal response to a call (the number of antiphonal calls each bat made divided by the number of calls each bat received). We used binomial tests to estimate a 95% confidence interval (CI) around these frequencies.

We used a permutation test to determine if bats made antiphonal calls more often than expected under a null model. We generated 5000 datasets expected from two bats calling independently of each other. To do this, we constructed fake sessions by combining the full session of call onset times from one random caller with that of a second random caller-session. This procedure preserves the timing of calling within bat and session, but removes any statistical relationship between the call times of the caller and receiver. We sampled these caller-session vocal sequences without replacement, and we removed any session in which a bat was randomly paired with itself. We measured the overall frequency of antiphonal calls in each permuted dataset and calculated 95% quantiles of the expected values under the null hypothesis.

## RESULTS

Dyadic contact calling rates were clearly predicted by allofeeding rates and by allogrooming rates. Under the caller-receiver model, an allofeeding rate that was one standard deviation higher corresponded to 1.7 times more contact calls in the recording session (95% Credible Interval (CrI) = [1.1, 2.8]), and an allogrooming rate that was one standard deviation higher corresponded to 2.4 times more contact calls (95% CrI = [1.3, 4.9]). However, greater genetic kinship did not clearly correspond to more contact calling (95% CrI = [0.6, 3.9], Fig. 1, Table S2). Evidence for higher calling rates among preferred cooperation partners, but not necessarily among kin, was also evident even from the social relations model (allofeeding incidence rate ratio (IRR): median = 1.73, 95% CrI = [1.08, 2.83]; allogrooming IRR: median = 2.40, 95% CrI = [1.26, 4.90]; kinship IRR: median = 1.49, 95% CrI = [0.58, 3.93], Fig. S2, S3; Table S2). The association between cooperative relationships and calling rates is therefore not fully explained by kinship.

**Figure 1.**
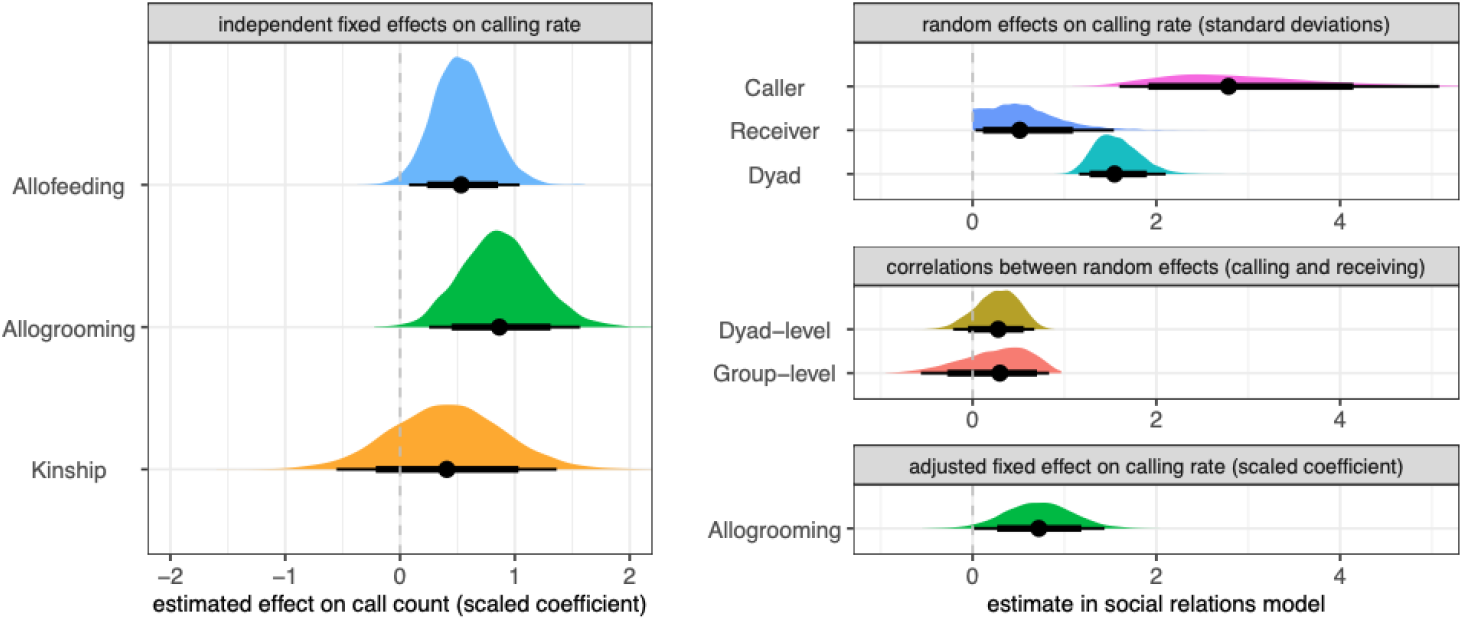
Contact calling rates are predicted by allofeeding and allogrooming. Left panel shows caller-receiver model coefficients for dyadic allofeeding, dyadic allogrooming, and kinship as independent predictors of contact calls with posterior distributions and means (with 80% and 95% Bayesian credible intervals). Right panels show results from social relations models with variances explained by caller, receiver, dyad (bounded at 0); correlations at the dyad and group levels (vocal reciprocity, bounded between -1 and 1); and the effect of allogrooming after adjusting for these other effects. In the social relations model, allogrooming estimates may be underestimated if preferred calling relationships (random effect of dyad) are not distinct from the relationship effect being measured by allogrooming. Social relations models with allofeeding (Fig. S2) and kinship (Fig. S3) had similar estimates. Full model results are shown in Table S2 and S3.

Some bats consistently called more than others and some bats consistently called more to certain partners. The clearest source of variation in calling rate was the identity of the caller, which explained more variation than the dyad and the receiver (Fig. 1). The effects of calls given on calls received was not clearly different from zero at either the group level or dyad level (Fig. 1, Table S2). For this reason, the highest estimate of calling reciprocity (from the social relations model with kinship: median correlation = 0.39; HDPI = [0.01, 0.72]) was far lower than our estimate of allogrooming reciprocity (median correlation = 0.81; HDPI = [0.62, 0.94], Fig. S3).

Only 1.87% (95% confidence interval = 1.46% to 2.36%) of calls made by common vampire bats were antiphonal (within 0.5 s of their partner’s calls, following Carter et al. (23)). This frequency was greater than expected from our null model (p < 0.0002, Fig. S4). Rates of antiphonal calling varied across bats (Fig. S5).

## DISCUSSION

### Contact calling is predicted by cooperative relationships in female vampire bats

Our results suggest that contact calls in common vampire bats are addressed to specific receivers and may be used to coordinate with preferred associates within groups. Contact calling among isolated pairs of adult female vampire bats was predicted by allogrooming and allofeeding rates, and these associations were not explained by kinship, which had weaker and uncertain effects on contact calling. Allogrooming was likely the best predictor of contact calling rates because it is relatively common; female vampire bats spend about 5% of their awake time allogrooming others (24). Although allofeeding is the most reliable indicator of willingness to help another individual, it is rarer and less precisely estimated than allogrooming.

Estimates of vocal reciprocity were far lower than allogrooming reciprocity for two reasons. First, common vampire bats rarely responded to each other’s calls (unlike white-winged vampire bats, which often respond to a group member’s contact calls within about one-third of a second (23)). Second, some common vampire bats were consistently far more vocal than others regardless of the receiver (Fig. 1). Dyadic contact-calling rates were therefore far more variable across callers compared to the weaker individual variation in allogrooming rates. Allogrooming relationships vary most at the dyad level and dyadic grooming is highly balanced and often mutual (Fig. S3).

### Adult female vampire bats have low rates of rapid responses

Antiphonal calling exchanges between adult group members occur in many group-living mammals, including Japanese macaques (25), common and pygmy marmosets (26–30), squirrel monkeys (31), Campbell’s monkeys (32), tamarins (33), male white-handed gibbons (34), mated pairs of Titi monkeys (4) and lemurs (3), sperm whales (35), beluga whales (36), killer whales (37), dolphins (38), elephants (39,40), meerkats (41), naked mole-rats (42), tree shrews (43), Spix’s disc-winged bats (44), and white-winged vampire bats (23), the closest living relative to common vampire bats. The frequency of this behavior across mammals may provide insights into the evolution of cooperative vocal turn-taking (45).

However, antiphonal exchanges in common vampire bats were rare. Group members rapidly exchange contact calls about ten times less than white-winged vampire bats (1.87% vs 18%, respectively (23)). It remains unclear why common vampire bats are less vocally responsive, but our results suggest yet-to-be-discovered differences in social behavior between these vampire bat species.

The function and pattern of antiphonal contact-calling relationships have been well studied in Spix’s disc-winged bats, *Thyroptera tricolor* (2,44,46), which form mixed-sex groups of related and closely associated bats that switch among leaf roosts on a nightly basis (47,48). To coordinate roost selection, group members produce contact calls called “inquiry calls” while in flight; roosting bats respond with distinctive “response calls” (44). Both inquiry and response calling rates are highly variable among callers, similar to common vampire bats (2). However, unlike vampire bats, contact calling in disc-winged bats is not predicted by association, kinship, or receiver, suggesting that disc-winged bats do not address contact calls to specific group members (2). Similarly, greater spear-nosed bats, *Phyllostomus hastatus*, use contact calls called “screech calls” to coordinate with group members, but they do not appear to use screech calls to recognize or address specific individuals (49–51).

### Implications for the vocal grooming hypothesis

The “vocal grooming” hypothesis, supported by studies of several primates (5,7,8) and other social mammals (38,53,54), posits that vocal exchanges contribute to social bonding in a similar fashion as allogrooming (55,56). One extension of this idea is that, over evolutionary time, the functions of physical grooming in primates were transferred to vocal grooming, and eventually replaced by spoken conversations in the human lineage (55,56). However, the more general idea of “vocal grooming” or “bonding at a distance” is that vocal communication plays a key role in forming and maintaining affiliative and cooperative relationships.

Our results are consistent with both a simple and a complex causal model of the vocal grooming idea. In the simpler causal model, vampire bats have a latent network of individualized social preferences, which then influence social behaviors such as co-roosting, clustering, allogrooming, allofeeding, and contact calling. According to this model, the so-called “effect” of allogrooming on contact calling is actually due to both behaviors (“physical grooming” and “vocal grooming”) being caused by the same underlying affiliative preference (or “social bond”). That is, changes in allogrooming and changes in contact calling are both caused by changes in this latent relationship, and a bat’s most preferred grooming recipient partner is also its most preferred recipient of feeding and calling. This scenario is best captured by the simpler caller-receiver model. The social relations model attempts to estimate the allogrooming relationship and the calling relationship as separate factors, but according to this simple model, these factors would be redundant measures of the same causal effect.

Alternatively, in the more complex causal model of “vocal grooming”, vampire bats have separate preferences for who they groom, feed, and call towards. According to this model, a bat’s most preferred grooming recipient could be its least preferred calling recipient. This scenario is better captured by the social relations model, which estimates a contact-calling relationship (dyad effect) as a separate source of variation than the social preference estimated by allogrooming. However, the greater complexity of this causal model means that the estimates of the effect sizes are less certain and harder to interpret due to confounding. For example, if there are strong cyclical feedbacks between and within interaction types (e.g. allogrooming A to B causes contact calling B to A which causes contact calling A to B), then estimating these causal effects would require future work that either (1) applies causal inference to larger datasets or (2) experimentally manipulates one interaction type (contact calling) and measures responses in another (allogrooming).

Under either model, our findings contribute to increasing evidence (14) that vocal interactions play a previously-underappreciated key role in how vampire bats form and maintain cooperative relationships.

## Ethics

Procedures with animals were approved by the Institutional Animal Care and Use Committee at The Ohio State University (protocol 2021A00000067).

## Data accessibility

Data and R code for reproducing analyses is available on GitHub (57).

## Declaration of AI use

We have not used AI-assisted technologies in creating this article.

## Authors’ contributions

Julia Vrtilek: data curation, formal analysis, investigation, methodology, project administration, visualization, writing—original draft, and writing—review and editing. Haley Gmutza: data curation, investigation, and writing—review and editing. Sydney Decker: data curation, investigation, and writing—review and editing. Gerald Carter: conceptualization, data curation, formal analysis, funding acquisition, investigation, methodology, project administration, resources, supervision, visualization, writing—original draft, and writing—review and editing. All authors gave final approval for publication and agreed to be held accountable for the work performed therein.

## Acknowledgements

We thank Tobias Nguyen for feedback that improved the manuscript.

## SUPPLEMENT

**Figure S1.**
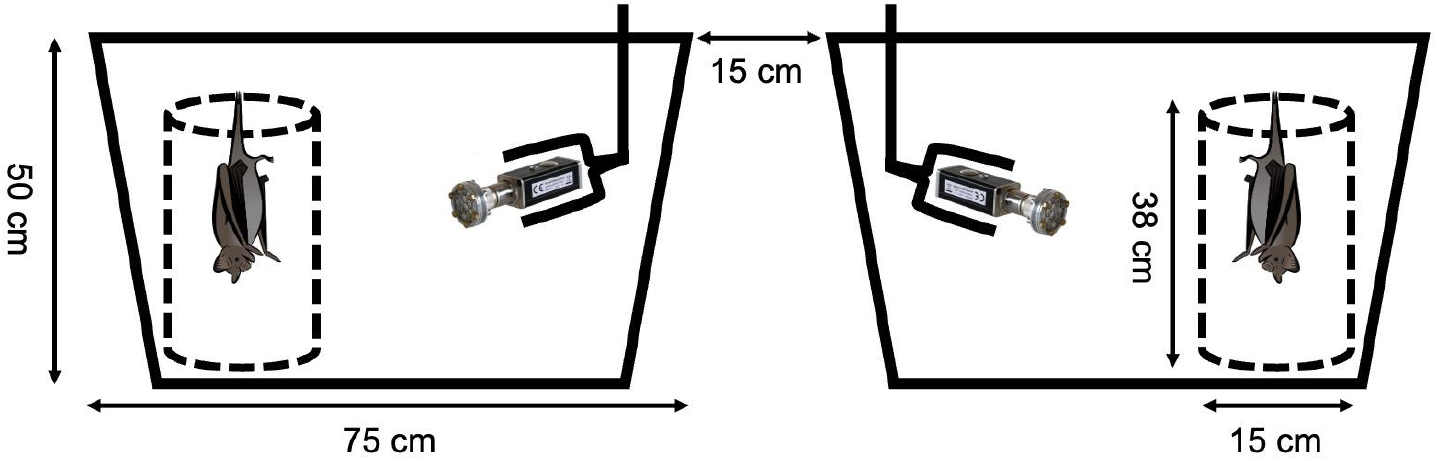
Setup for recording dyadic contact calling. Pairs of bats were physically and acoustically isolated from the rest of the colony, then recorded in adjacent plastic bins lined with acoustic foam. Within each bin, the bat was placed in a plastic mesh tube to keep it in front of the microphone, about 10-30cm away. Not to scale.

**Figure S2.**
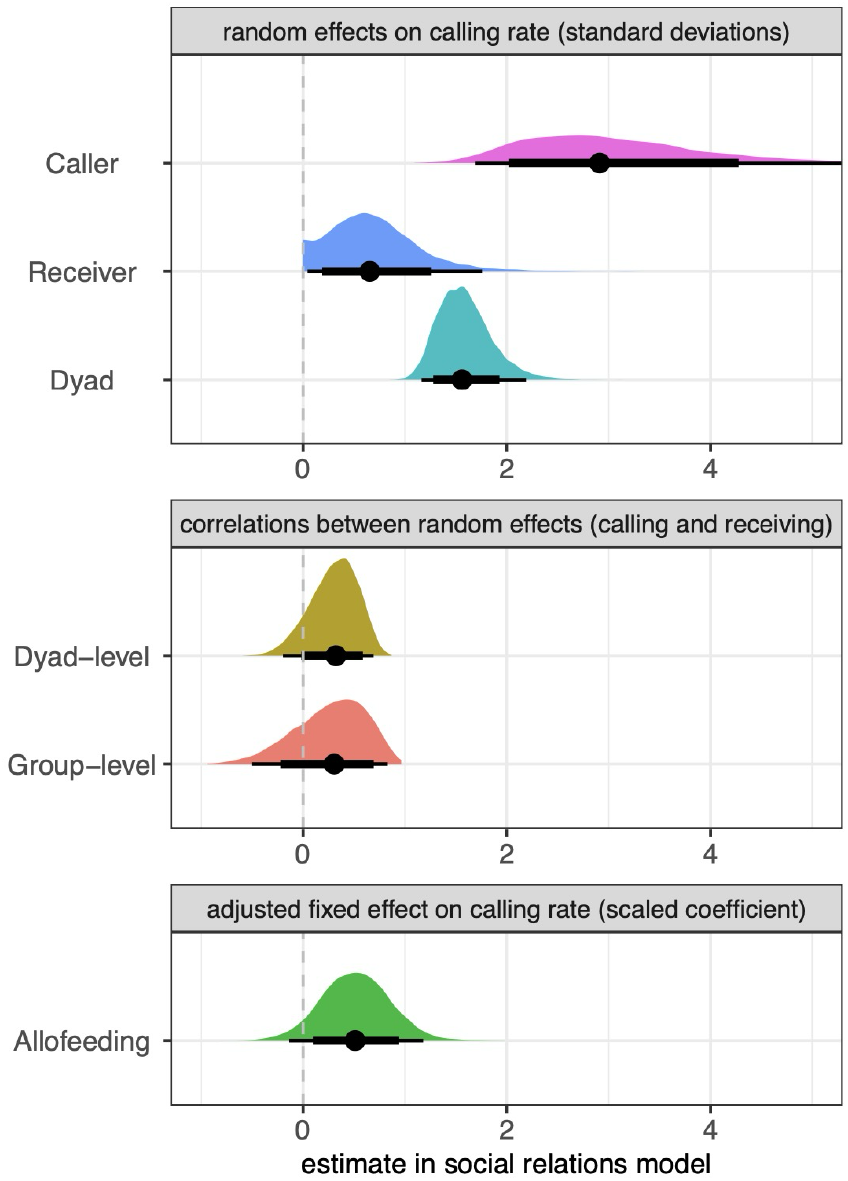
Estimates from social relations model estimates with allofeeding as measure of social preference. Posterior probability distributions and means (with 80% and 95% Bayesian credible intervals) are shown for sources of variances explained by caller, receiver, and dyad, correlations at the dyad and group levels, and the effect of allofeeding after adjusting for other effects. Note that allofeeding estimates may be underestimated because these models assume that preferred calling relationships (random effect of dyad in top panel) are distinct from the relationship effect being measured by allofeeding (bottom panel).

**Figure S3.**
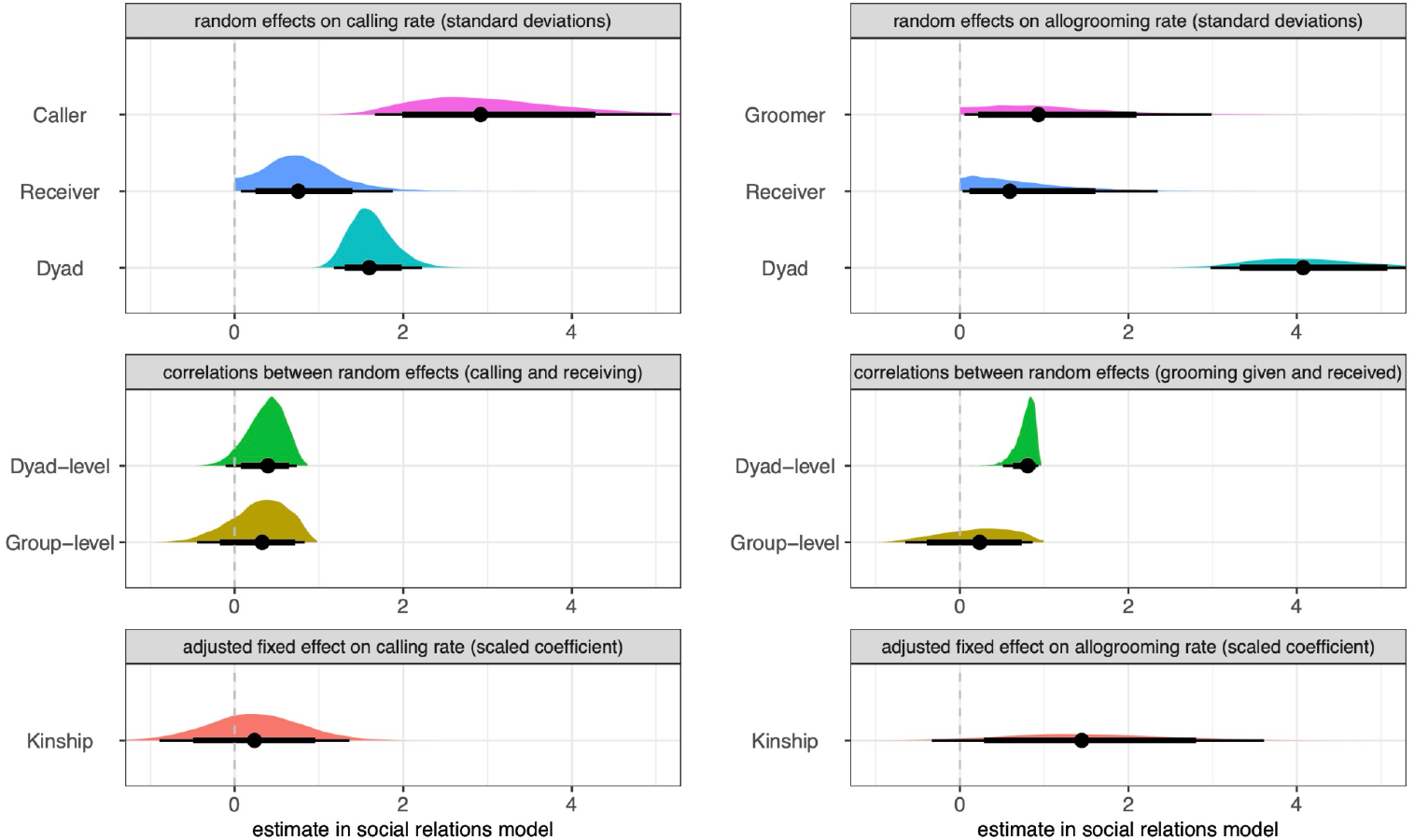
Contact calling is less reciprocal than allogrooming. Plots show a comparison of social relations model estimates with kinship predicting calling rate (left side panels) or allogrooming (right panels) using the same bats. Compared to contact calling which varies across callers, allogrooming varies across dyads and is highly reciprocal within dyads. Kinship does not clearly predict calling or allogrooming.

**Figure S4.**
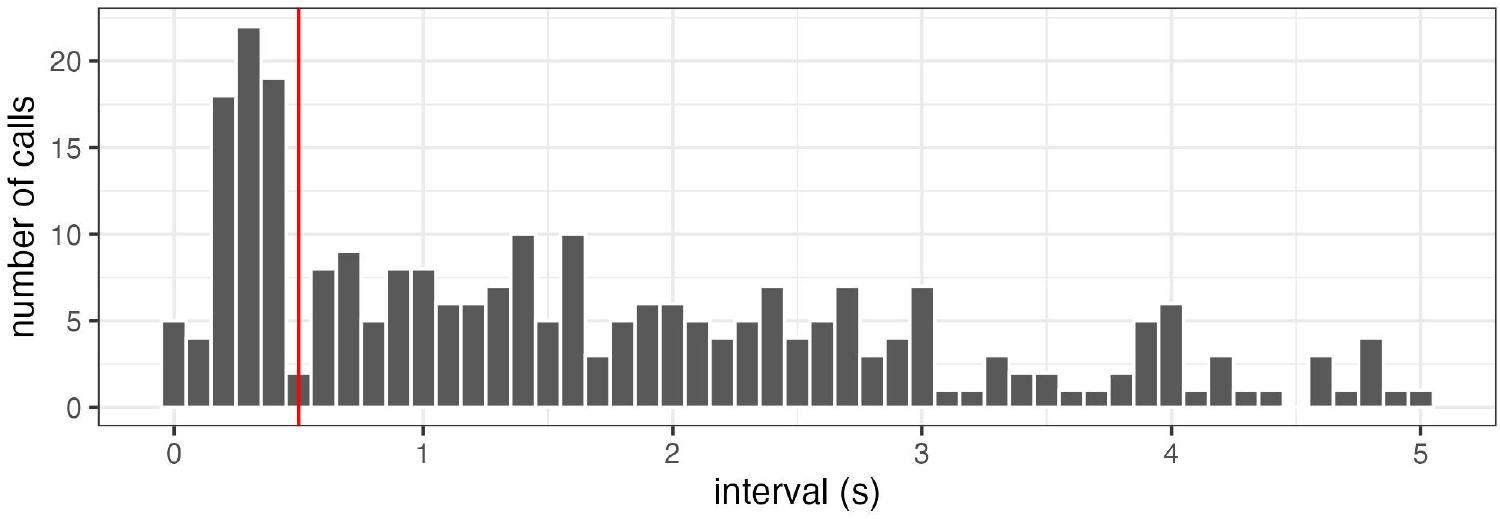
Distribution of latency between contact calls from different bats. Inter-call intervals are shown for 5 seconds after a call. Red line indicates a latency of 500 ms, the threshold for defining an antiphonal response, following Carter et al. (23).

**Figure S5.**
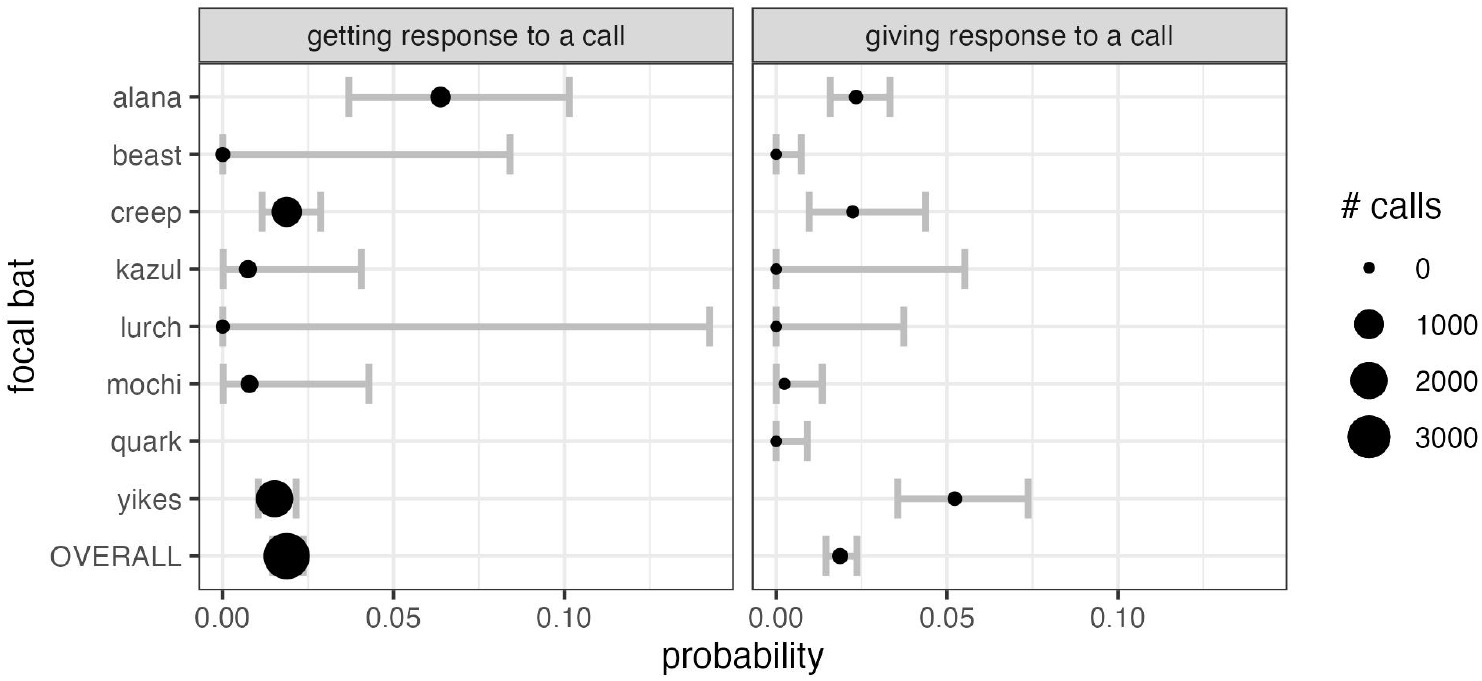
The probability of each bat getting a response to a call (panel 1) and giving a response to a call (panel 2). Error bars show 95% CIs from a binomial test.

**Table S1.**
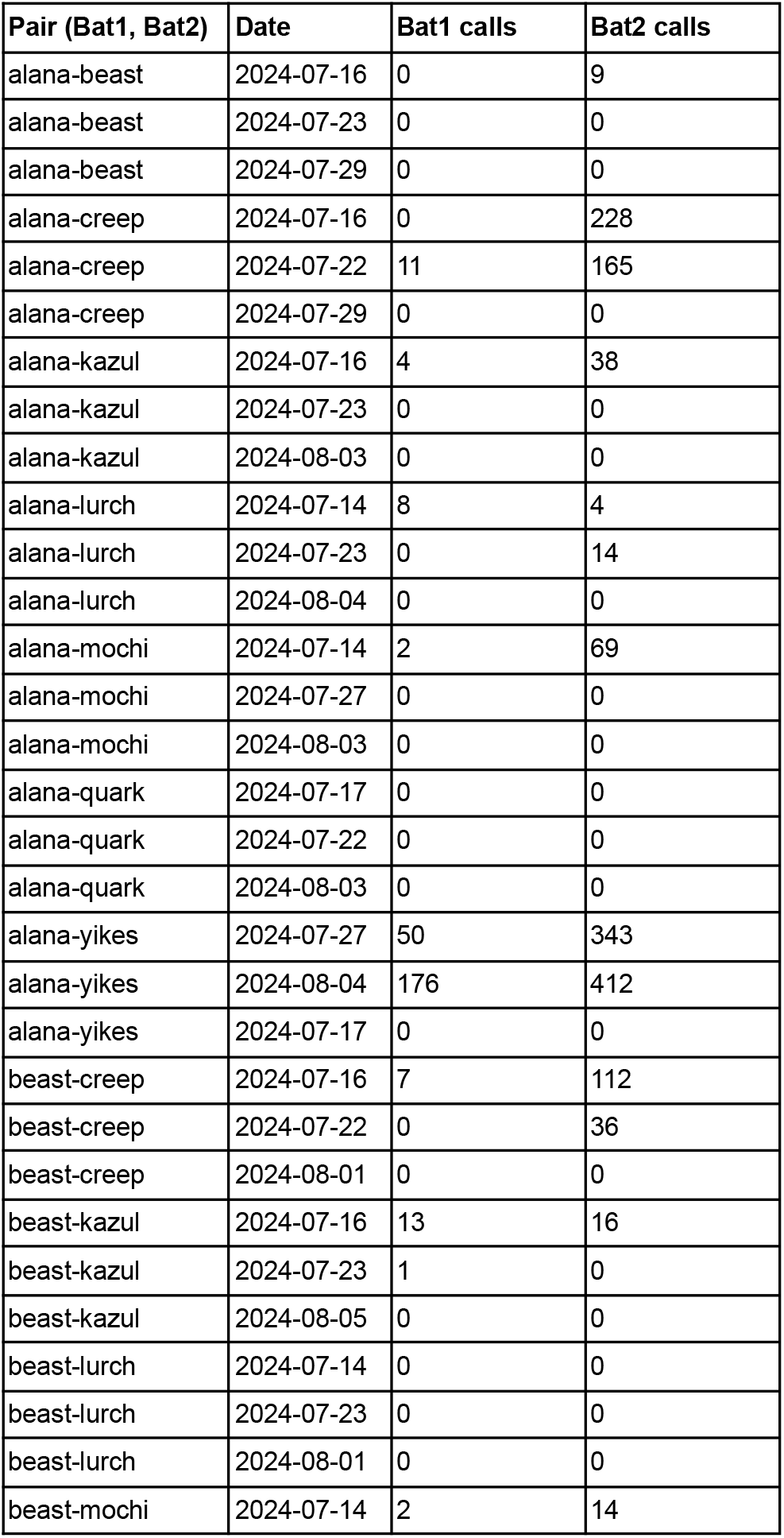
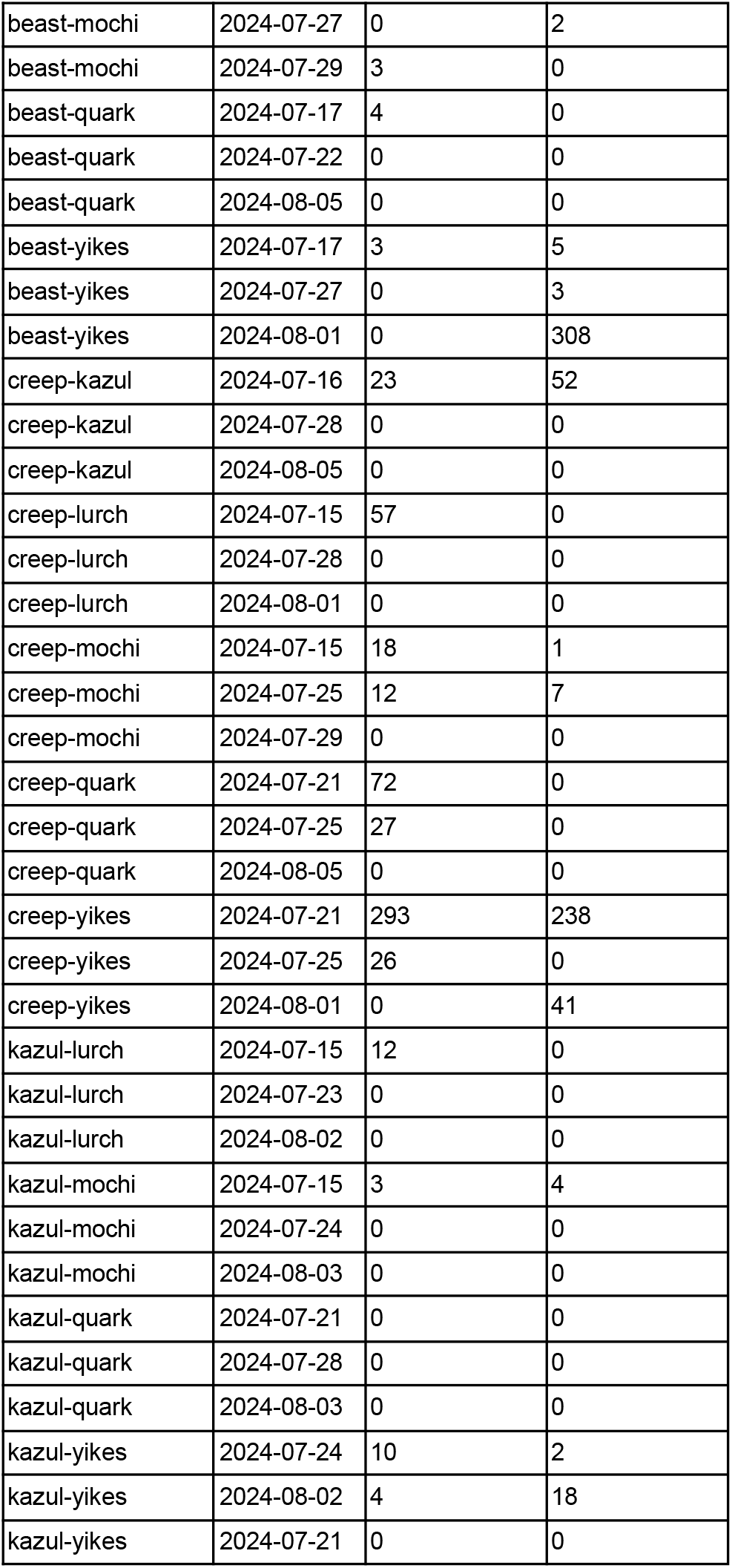
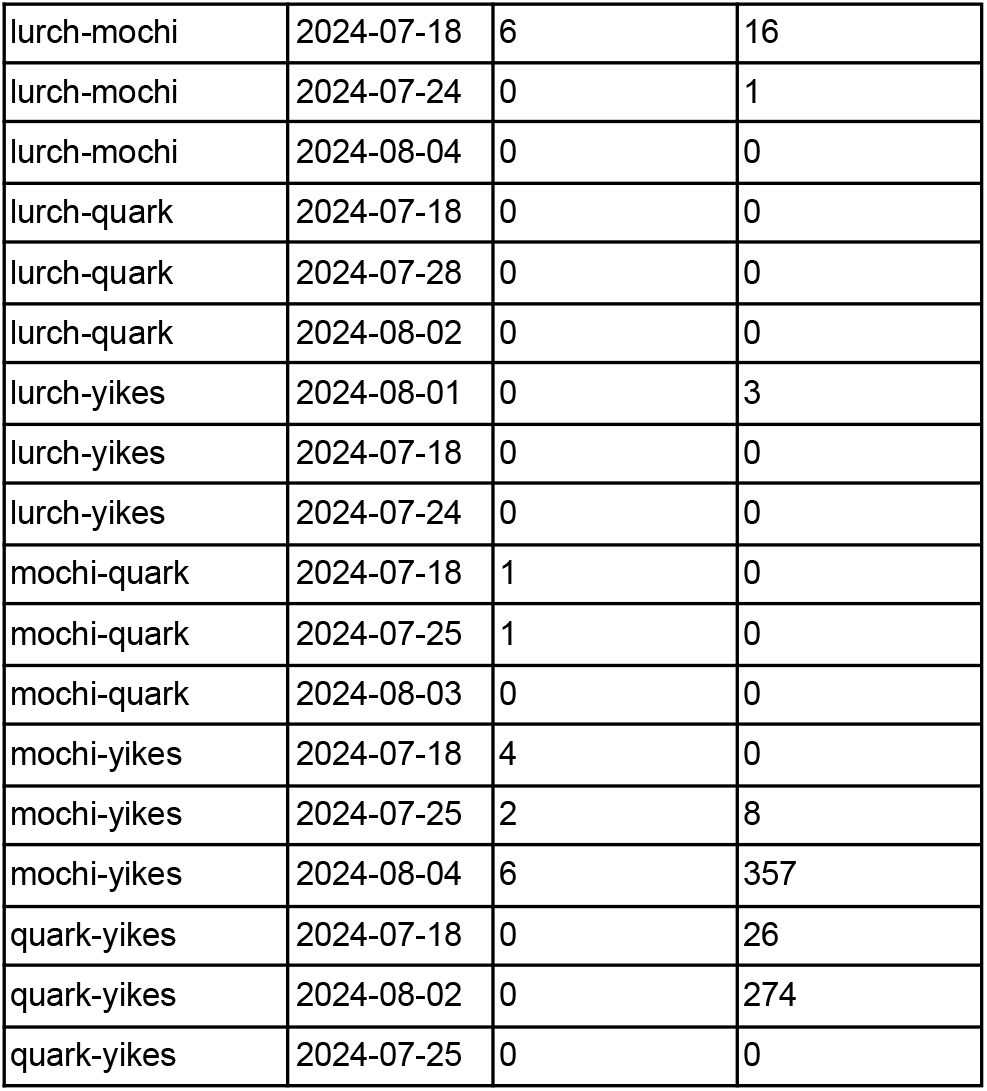
Contact call counts per bat pair and recording session.

**Table S2.** Model estimates for predictors of call count. Results from two models addressing different questions, each run with 3 different social predictors (allofeeding, allogrooming, or kinship). The caller-receiver model considers three causes of calling rate: the caller, the receiver, and the listed social predictor. The social relations model considers all of the above, but also assumes a) that bats can have preferred calling partners that are not fully explained by social preferences (estimated with listed predictor) and b) that random effects of caller and receiver are correlated at both the group- and dyad-level. For each model, we list the estimated effect of each predictor, with lower and upper bounds of a 95% CI, and the proportional increase in contact calling for each standard deviation increase in predictor (IRR) with lower and upper bounds.

| Type | Predictor | Estimate | Est. Lower<br>95% CrI | Est. Upper<br>95% CrI | IRR | IRR<br>Lower | IRR<br>Upper |
| --- | --- | --- | --- | --- | --- | --- | --- |
| Caller-Receiver<br>Model | Allofeeding | 0.55 | 0.07 | 1.04 | 1.73 | 1.08 | 2.83 |
| Caller-Receiver<br>Model | Allogrooming | 0.88 | 0.23 | 1.59 | 2.40 | 1.26 | 4.90 |
| Caller-Receiver<br>Model | Kinship | 0.40 | -0.55 | 1.37 | 1.49 | 0.58 | 3.93 |
| Social Relations<br>Model | Allofeeding | 0.52 | -0.14 | 1.18 | 1.68 | 0.87 | 3.25 |
| Social Relations<br>Model | Allogrooming | 0.72 | 0.01 | 1.44 | 2.06 | 1.01 | 4.20 |
| Social Relations<br>Model | Kinship | 0.24 | -0.89 | 1.36 | 1.26 | 0.41 | 3.90 |

**Table S3.**
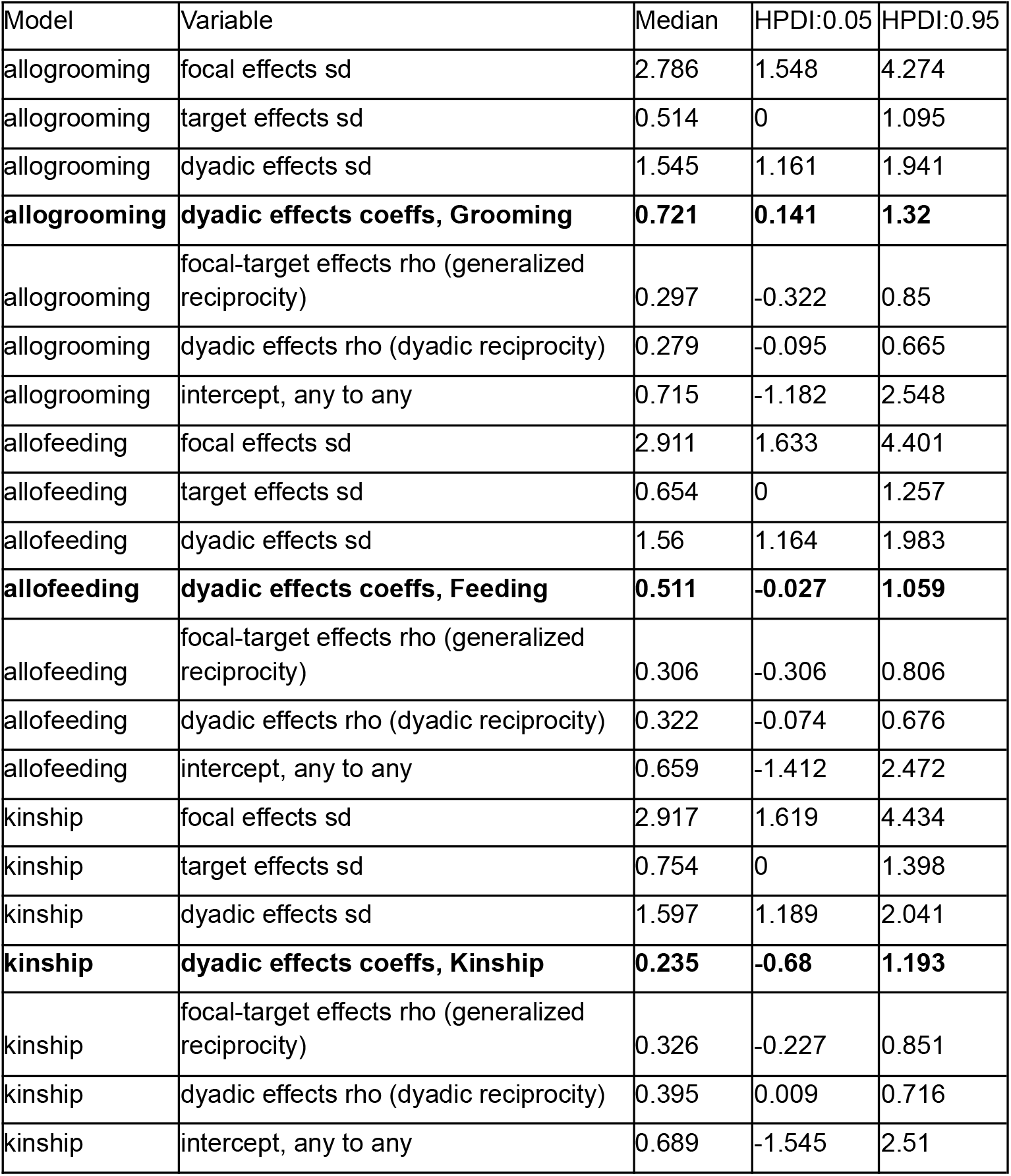
Social relations model estimates. For each social predictor (allogrooming, allofeeding, or kinship), we list the posterior median and highest 90% posterior density interval for the following: standard deviation for effect of caller (focal), receiver (target), and caller-receiver dyad; dyadic effect of the social predictor; estimate for generalized reciprocity (calls given to and received from *any bat*); estimate for dyadic reciprocity (calls given and received *within pair*); and the intercept of calls from and to any bat.

